# *En route* activity of Chilean Elaenia, a long-distance migratory bird in South America

**DOI:** 10.1101/2024.08.26.609733

**Authors:** Victor R. Cueto, Cristian A. Gorosito, Geoffrey Brown, Alex E. Jahn

## Abstract

The logistics of measuring activities that occur at fine temporal scales, such as short stopovers that last a few hours, has proven very challenging when studying small migratory birds. Here, we deployed multi-loggers equipped with an accelerometer and thermometer on Chilean Elaenia (*Elaenia chilensis*) to evaluate their activity patterns while they undertook their annual migration from their Patagonian breeding grounds to non-breeding zones in Brazil. Results show that elaenias can fly at altitudes of >1500 masl and migrate nocturnally, providing the first evidence of this behavior in a Neotropical austral migrant. Although most migration flights lasted less than 8 h, one individual flew non-stop for more than 28 h. Overall activity patterns (e.g., flight and stopover duration) were not substantially different between pre- and post-breeding migration. This technology offers a window into the migratory behavior of small birds that migrate within the Neotropics at a finer temporal scale than previously possible.

## 1. Introduction

Migration allows birds to improve their fitness by accessing greater seasonal resources, while avoiding other limitations, such as adverse weather conditions, competition for resources or parasitism (Alerstam et al. 2003, Scott 2010). This benefit must outweigh the costs associated with these movements, such as a greater risk of mortality (e.g., predation), time restrictions in the annual cycle (e.g., less time for reproduction), and greater expenditure of metabolic energy (Alerstam et al. 2003). Migrants accomplish this by utilizing a suite of migratory strategies, such as nocturnal migration (Alerstam 2011), long-distance flights or alternatively, several flights with intermediate stops (Scott 2010), and flying at different altitudes (reviewed by Newton 2023).

Although past and current research on bird migration focuses on species that breed in North America and Europe, recent progress has been made in describing basic migration patterns for species that breed and migrate at tropical and south-temperate latitudes in South America (reviewed by Jahn et al. 2020). Since South America is more seasonally buffered and characterized by lower seasonal predictability between years than North America (Lisovski et al. 2017), there should be a lower selective pressure for birds that breed within the Neotropics to accurately time migration relative to those that breed at north-temperate latitudes (Jahn et al. 2020). Thus, Neotropical austral migratory birds (*sensu* Cueto and Jahn 2008), which migrate wholly within South America, should migrate slower than ecologically similar species breeding at north-temperate latitudes, exhibiting an overall energy-rather than time-selected migration strategy (Jahn and Cueto 2012, Jahn et al. 2020). Such a slower migratory strategy would be characterized by, for example, by longer and more frequent stopovers and a steeper rise in cumulative flight hours over time (e.g., Backmän et al. 2017), yet our knowledge of *en route* activity of small migratory birds on an hourly or even daily basis remains very poorly understood. For example, substantial evidence exists to show that passerines generally migrate faster during pre-vs. post-breeding migration (reviewed by McKinnon et al. 2013, Nilsson et al. 2013), yet for most migratory bird species across the planet, we still have little to no information on the hourly changes in activity (e.g., overnight stopover duration) from which seasonal patterns of movement rate and timing arise.

The aim of our study was to analyze *en route* activity of a Neotropical austral migrant, the Chilean Elaenia (*Elaenia chilensis*) at an hourly scale. Elaenias spend the non-breeding season >6000 km from their Patagonian breeding sites, carrying out one of the longest known migrations among Neotropical austral migrant passerines (Bravo et al. 2017, Jara et al. 2024), making them an ideal model to study *en route* activity during long-distance migration in the Southern Hemisphere. We describe elaenias activity patterns (e.g., diurnal vs nocturnal migration, stopover duration) using accelerometers, which have proven to be a powerful tool for resolving migratory activity at fine temporal scales in studies based in the Northern Hemisphere (e.g., Briedis et al. 2020, Macías-Torres et al. 2022, Vīgants et al. 2023, Adamíck et al. 2024).

## 2. Material & Methods

### 2.1. Study site

We conducted our research at El Principio Ranch (42°5649S, 71°2330W, 566 m.a.s.l.), in the NW of Chubut province, Argentina. The vegetation corresponds to the Valdivian Forest Province of the Andean Region (Morrone 2001). The forest in the ranch is mainly composed of *Maytenus boaria, Schinus patagonicus, Nothofagus antarctica*, and *Austrocedrus chilensis* trees, and the understory is dominated by *Berberis microphylla* shrubs. This forest is common in valleys and slope of hills in the eastern portion of the Patagonian Forest in Chubut Province, and forms part of the ecotone between the forest and the steppe (Kitzberger 2012). The climate is characterized by cold and wet winters and mild but dry summers. The rainy season encompasses the months of April–September (fall–winter), when 68% of the 704 mm of annual precipitation falls as rain and snow. The remainder of the year constitutes the dry season (spring–summer). All climate data are from the Río Percey meteorological station (period 1998–2017, Hidroeléctrica Futaleufú S.A.), located 9 km NW of our study area.

### 2.2. Study species

Taxonomy of the Chilean Elaenia is complex and the species-level classification remains contentious (Pearman and Areta 2020). Therefore, we have adopted the taxonomy of the IOC World Bird List (Gill et al. 2024). *Elaenia chilensis* is the same species as *Elaenia albiceps chilensis* of Billerman et al. (2022) and Clements et al. (2023).

Chilean Elaenia (Supplementary material Figure S1) is a long-distance Neotropical austral migrant which exhibits a loop migration pattern, using a single pre-breeding migration route and three different post-breeding migration routes (Bravo et al. 2017, Supplementary material Figure S1). Elaenias move quickly through desert and grassland-dominated regions during post-breeding migration (Bravo et al. 2017), at times stopping over for extended periods (Bravo et al. 2017). During pre-breeding migration, they make a one-way trip of more than 5500 km between their non-breeding grounds in northern Brazil and breeding zones in western Patagonia (Bravo et al. 2017, Jara et al. 2024). Elaenias migrate at similar rates to those of ecologically similar species that breed at north-temperate latitudes (Bravo et al. 2017). During the breeding period, it is the most abundant bird species in the Patagonian Forest (Cueto and Gorosito 2018), where it contributes to forest regeneration (Bravo et al. 2015). Males begin to arrive at our study site in mid-October (Bravo et al. 2017, Cueto and Gorosito 2018), while females begin to arrive in early November (Cueto and Gorosito 2018, Gorosito 2020). Adults begin post-breeding migration between mid- and late February (Bravo et al. 2017, Cueto and Gorosito 2018), although occasionally some of them stay at our study site until early March (Cueto and Gorosito 2018, Gorosito 2020).

### 2.3. Field work

We captured elaenias as part of a systematic bird monitoring program at El Principio Ranch during the October-March 2021–2022 breeding season (i.e., spring-summer) using mist nets (3 × 12 m, 38-mm mesh size) set in two groups of 10 nets each, located 200 m apart, and nets within each group 70 - 100 m apart. We alternated the operation of each group every 10 days, operating nets for 5 h, starting at sunrise. We avoided mist-netting during rainy or windy days. Captured birds were banded with numbered metal bands and a unique combination of up to three Darvic color bands. For each captured individual, we measured unflattened wing chord (from the carpal joint to tip of the longest primary), tail length and body mass. All individuals were sexed as described in Cueto et al. (2015).

Twenty adult elaenias (2 females and 18 males) were fitted with a BitTag, a multi-sensor logger that comprises a real-time clock, a microcontroller with large non-volatile memory and built in temperature sensor, and an accelerometer (Brown et al. 2023). The accelerometer measures acceleration in three dimensions at a ± 4 g range at a sampling rate of 50 Hz (12 Hz when bird is inactive). Changes in acceleration allow identifying periods when an animal is active or inactive, with an animal being considered to be inactive when there have been no significant changes in acceleration in the preceding 0.5 seconds. BitTags aggregate activity data by counting active seconds during the aggregation period. We used a threshold for active/inactive transitions of 0.5 g, with an aggregation period of 5 min. We used a BitTag weighing 0.6 g mass (Brown et al. 2023) to allow for deployment on elaenias, but which resulted in the use of a smaller battery, restricting the available energy for running the accelerometer throughout the year. Due to this restriction, BitTags were pre-programmed on a schedule that defined the dates when the accelerometer collected data. The reduction of measurement periods substantially reduced the amount of data collected, but prolonged tag operation time by minimizing power consumption. In our study, we ran two measurement periods and one hibernation period. Measurement periods ran from February 15 to April 21 and September 11 to November 20, with hibernation periods from April 22 to September 10. Timing of measurement sequences was selected to match periods of migration previously identified by light-level geolocators (Bravo et al. 2017).

BitTags were attached using a leg-loop harness (Rappole and Tipton 1991) made of StretchMagic (0.7 mm diameter, Pepperell Braiding Co.). The combined mass of the tag and harness was 0.7 g and never exceeded 5% of the mass of elaenias (mean ± SE = 14.8 ± 0.5 g, n = 20). All birds flew well upon release and we tried to recapture them in the following breeding seasons to recover the tags, both through the aforementioned systematic sampling with mist nets and by searching their territories throughout the study area. Once a tagged elaenia was found, we placed a 6-m-long mist net within the defended territory and attracted the individual using a ceramic model of an elaenia and a speaker placed nearby that emitted the species’ vocalizations. Recaptured individuals showed no sign of injury from the BitTag or harness. During the study, the average return rate of elaenias without BitTags was 46% and 32% for males and females, respectively. The return rate for the 18 males we tagged was 27%, and we did not recapture either of the two tagged females.

### 2.4. Data analysis

Flight periods based on the accelerometer data were identified when the BitTag recorded more than 50% of activity during each aggregation period of 5 min. Using the previously-defined flight period, the start and end points of flight periods, and hence flight duration, were calculated to a 5 min resolution, provided that activity was maintained >50% and all recorded in sequence. The end of a flight was defined as when activity fell below 50%. Periods between well-defined flight periods were recorded as stopovers or breeding site residency. In some cases, BitTags recorded short flight periods (less than 1 h, usually with a duration between 5 min and 30 min) during periods of low activity (i.e., <50%). In such cases, we considered that the bird had landed and considered the full sequence as a stopover period when calculating its duration. Differences on fight and stopover activity between pre- and post-breeding migration were statistically evaluated using Student’s t test.

BitTags recorded temperature every 35 min, and we used the last temperature recorded before departure and the first temperature recorded during the first pre-or post-breeding migration flight. The temperature recorded by the BitTag is influenced by the bird’s body temperature (since the difference between the temperature measured by the BitTag and ambient temperature measured by a local meteorological station at the initiation of the first post-breeding migration flight was 18.8 ± 6.0 °C, Supplementary material Table S1). However, we consider that the absolute difference between the temperature measured by the BitTag prior to the first pre- or post-breeding migration flight and when the bird was on migration is a reliable estimate of flight altitude, since atmospheric temperature reduces in a predictable way, with an average drop of 0.7 °C with every 100 m height (Ahrens 2009).

## 3. Results

We recaptured five tagged elaenias, all of them males. Four of them recorded activity throughout pre- and post-breeding migration, while one stopped recording on 23 February (i.e., 8 days after the start date).

During both pre- and post-breeding migration, all elaenias During both pre- and post-breeding migration, all elaenias flew primarily during nocturnal hours, departing in the evening and stopping the next morning (Figure 1). During pre-breeding migration, they flew on average 6.8 ± 1.0 h (mean ± SE) per night (range: 1.2 - 11.5 h), totaling 99.1 ± 5.3 h of flight across 14.8 ± 2.1 nights (Table 1, Figure 1) to complete the journey from the non-breeding area to the Patagonian breeding site. When elaenias initiated the first pre-breeding flight, they flew at an altitude of 235. 7 ± 142.6 m (range: 100.0 - 385.7 m, Table 2).

**Table 1.** Migratory flight activity of Chilean Elaenias tagged with BitTags in the forest-steppe ecotone at El Principio Ranch, Chubut Province, Argentina. SD = standard deviation.

| BitTag<br>ID | First flight | Last Flight | Number<br>of flights | Total duration<br>(h) | Mean duration<br>(h) | SD<br>(h) | Range<br>(h) |
| --- | --- | --- | --- | --- | --- | --- | --- |
| Pre-breeding migration |  |  |  |  |  |  |  |
| DD | September 19 | October 31 | 13 | 97.9 | 7.5 | 3.4 | 2.0-10.3 |
| EA | September 19 | October 30 | 15 | 106.7 | 7.1 | 3.3 | 1.6- 11.5 |
| E5 | September 21 | November 05 | 18 | 97.4 | 5.4 | 3.4 | 1.2-10.1 |
| E7 | October 10 | November 17 | 13 | 94.5 | 7.3 | 2.8 | 1.8-10.2 |
| Post-breeding migration |  |  |  |  |  |  |  |
| DD | February 16 | April 20 | 19 | 182.2 | 9.6 | 6.0 | 2.8-28.9 |
| EA | February 19 | April 13 | 14 | 106.0 | 7.6 | 3.5 | 1.2-11.3 |
| E5 | March 03 | April 16 | 17 | 135.0 | 7.9 | 3.3 | 4.5-11.4 |
| E7 | March 03 | April 13 | 17 | 136.3 | 8.0 | 3.4 | 1.5-10.9 |

**Table 2.** Date, time, the last temperature recorded by BitTags before the first pre-breeding migration flight (Tpre), the first temperature recorded during that flight (Tmig), the difference between the two values (Tpre-Tmig), and estimated flight altitude during the first pre-breeding migration flight of Chilean elaenias tagged with BitTags in the forest-steppe ecotone at El Principio Ranch, Chubut Province, Argentina. Altitude above sea level is not calculated because departure locality in the non-breeding area is unknown; SD = standard deviation.

| BitTag ID | Date | Pre-migration |  | On-migration |  | Tpre-Tmig (°C) | Altitude (m) |
| --- | --- | --- | --- | --- | --- | --- | --- |
|  |  | Time | Tpre (°C) | Time | Tmig (°C) |  |  |
| DD | September 19 | 1725 | 35.9 | 1800 | 33.2 | 2.7 | 385.7 |
| EA | September 19 | 2350 | 33.1 | 2425 | 32.2 | 0.9 | 128.6 |
| E5 | September 21 | 1750 | 36.8 | 1825 | 36.1 | 0.7 | 100.0 |
| E7 | October 10 | 2055 | 34.2 | 2130 | 31.9 | 2.3 | 328.6 |
| Mean |  |  | 35.0 |  | 33.4 | 1.7 | 235.7 |
| SD |  |  | 1.7 |  | 1.9 | 1.0 | 142.6 |

**Figure 1.**
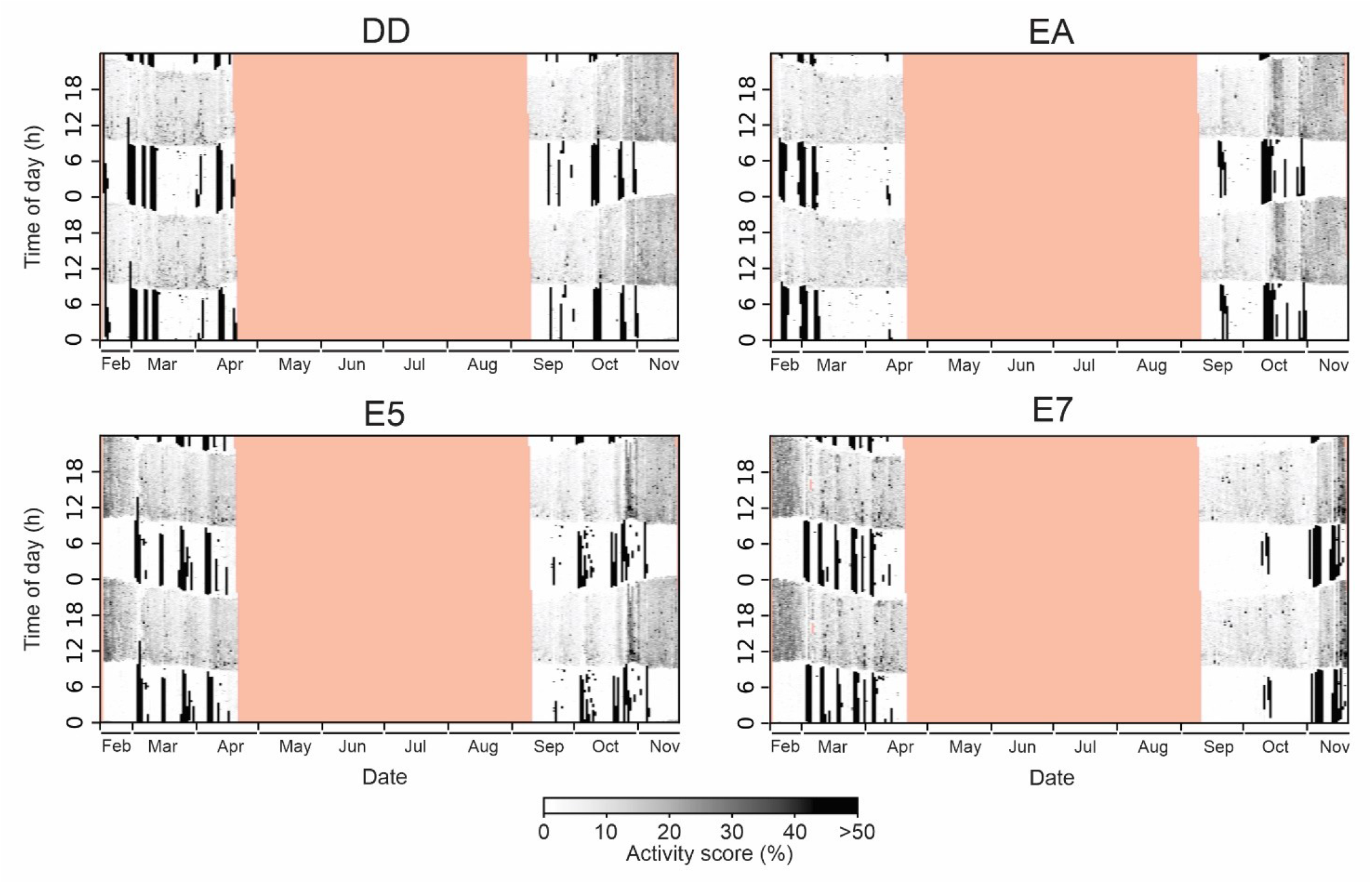
Actograms depicting activity of Chilean Elaenias tagged with BitTags in the forest-steppe ecotone at El Principio Ranch (Chubut Province, Argentina) during pre-breeding migration (September-November) and post-breeding migration (February-April). The amount of activity measured in 5 min periods increases with darker shading (black areas represent continuous flapping flight during >50% of every 5 min, and white areas represent the absence of flapping flight (0% of every 5 min). Pink areas indicate dates that accelerometers did not record activity. Letters above each actogram refer to BitTag ID.

During post-breeding migration, elaenias flew for on average 8.3 ± 0.9 h per night (range: 1.2 - 28.9 h), totaling 139.9 ± 31.5 h of flight during 16.8 ± 2.1 nights (Table 1, Figure 1) throughout the journey from the breeding site to their non-breeding areas in Brazil. When elaenias began the first post-breeding migration flight, they flew at an altitude of 1225.7 ± 634.7 m (range 771.4 - 2185.7 m, Table 3), corresponding to 1791.7 meters above sea level. Mean flight time, total flight time and number of flight nights were statistically non-significant between pre- and post-breeding migration (t = 2.21, P = 0.07; t = 2.51, P = 0.08; t = 1.28, P = 0.25, respectively).

**Table 3.** Date, time, the last temperature recorded by BitTags before the first post-breeding migration flight (Tpre), the first temperature recorded during that flight (Tmig), the difference between the two values (Tpre-Tmig), and estimated flight altitude during the first post-breeding migration flight of Chilean elaenias tagged with BitTags in the forest-steppe ecotone at El Principio Ranch, Chubut Province, Argentina. The study site’s altitude above sea level is 566 m; SD = standard deviation.

| BitTag ID | Date | Pre-migration |  | On-migration |  |  | Flight altitude (m) | Flight altitude above sea level (m) |
| --- | --- | --- | --- | --- | --- | --- | --- | --- |
|  |  | Time | Tpre (°C) | Time | Tmig (°C) | Tpre-Tmig (°C) |  |  |
| DD | February 16 | 1900 | 39.2 | 1935 | 33.8 | 5.4 | 771.4 | 1337.4 |
| EO | February 16 | 1935 | 37.9 | 2010 | 26.9 | 11.0 | 1571.4 | 2137.4 |
| EA | February 19 | 1845 | 39.9 | 1920 | 34.5 | 5.4 | 771.4 | 1337.4 |
| E5 | March 03 | 1855 | 37.1 | 1930 | 31.3 | 5.8 | 828.6 | 1394.6 |
| E7 | March 03 | 1855 | 36.8 | 1930 | 21.5 | 15.3 | 2185.7 | 2751.7 |
| Mean |  |  | 38.2 |  | 29.6 | 8.6 | 1225.7 | 1791.7 |
| SD |  |  | 1.3 |  | 5.4 | 4.4 | 634.7 | 634.7 |

During pre-breeding migration mean (± SD) number of stopover days was 13.8 ± 2.4 days (range: 12 - 17 days, Table 4, Figure 1), with a mean stopover duration of 68.4 ± 5.0 h (range: 13.4 - 472.8 h, Table 4), and a total stopover duration of 939.5 ± 168.8 h. During post-breeding migration, the mean number of stopover days was 15.8 ± 2.1 days (range: 13 - 18 days, Table 4, Figure 1) with a mean stopover duration of 68.6 ± 17.3 h (range: 2.5 - 811.4 h, Table 4), and a total stopover duration of 1067.6 ± 230.4 h. Number of stopover days, mean stopover duration and total stopover duration were statistically non-significant between pre- and post-breeding migration (t = 1.28, P = 0.25; t = 0.02, P = 0.98; t = 0.90, P = 0.40, respectively).

**Table 4.** Migratory stopover patterns of Chilean elaenias tagged with BitTags in the forest-steppe ecotone at El Principio Ranch, Chubut Province, Argentina. SD = standard deviation.

| BitTag ID | First Stop | Last Stop | Number of days | Total duration (h) | Mean duration (h) | SD (h) | Range (h) |
| --- | --- | --- | --- | --- | --- | --- | --- |
| Pre-breeding migration |  |  |  |  |  |  |  |
| DD | September 20 | October 31 | 12 | 895.4 | 74.6 | 81.6 | 13.5-109.4 |
| EA | September 20 | October 30 | 14 | 880.0 | 62.9 | 111.6 | 13.4-424.9 |
| E5 | September 21 | November 05 | 17 | 1184.4 | 69.7 | 118.8 | 15.0-451.3 |
| E7 | October 11 | November 16 | 12 | 798.1 | 66.5 | 130.9 | 13.9-472.8 |
| Post-breeding migration |  |  |  |  |  |  |  |
| DD | February 18 | April 02 | 18 | 1339.4 | 74.4 | 119.7 | 12.8-234.2 |
| EA | February 20 | April 13 | 13 | 1175.1 | 90.4 | 219.9 | 13.3-811.4 |
| E5 | March 04 | April 16 | 16 | 907.8 | 56.7 | 63.8 | 13.3-183.2 |
| E7 | March 04 | April 13 | 16 | 847.9 | 53.0 | 57.7 | 2.5-184.9 |

## 4. Discussion

Our study is the first to describe the activity patterns of a Neotropical austral migrant species *en route*, demonstrating that Chilean Elaenia migrates nocturnally, as has been shown in migratory passerines of the Northern Hemisphere (Berthold 2001, Newton 2023). Elaenias reached altitudes >1500 masl and, in general, their flights lasted 6 to 8 h per night, but individual DD flew more than 28 h non-stop during the first post-migration flight (see Figure 1). Overall, elaenia activity patterns during pre-breeding migration were not substantially different from that of post-breeding migration.

Migratory birds show great variability in the duration of stopovers, from a few hours to several weeks (Newton 2023). Elaenias exhibited two types of stopovers, one of a short duration and another of a substantially longer duration. Short duration stops were characterized by stops that started during the early morning, lasting ∼13 hours and initiating a new flight at dusk. Longer stopovers lasted several days and occurred intermittently during the entire migratory journey, during pre- and post-breeding migration. The first type of stopover is similar to that reported by Cueto et al. (2016) when elaenias were traveling during pre-breeding migration through the Monte Desert in west-central Argentina. Elaenias are primarily a forest edge and shrubland species, and during migration tend to cross unsuitable environments, such as grasslands and deserts (Bravo et al. 2017), executing prolonged stopovers after these flight periods. However, unlike birds in other migratory systems that must cross inhospitable environments, such as the Mediterranean Sea or the Gulf of Mexico without stopping (Newton 2023), elaenias executed multiple stopovers throughout the journey, which suggests that they experience fewer periods of sustained effort than birds that must cross high mountain ranges or oceans.

Although elaenias tended to travel faster during pre-than post-breeding migration, the differences were not substantial (11% fewer number of nights flying, 29% less total flight time and 24% less mean flight time). This could be due to the fact that during prebreeding migration, elaenias use only one route, while during post-breeding migration they use at least three different routes, some longer than others (Bravo et al. 2017). Additional evidence that elaenias are not under high selective pressure to arrive as early as possible to the breeding area is that they arrive at our study site more than a month before egg laying begins (Gorosito et al. 2022). Follow up research could thus test the prediction that elaenias have a migration strategy more associated with energy-rather than time-selection, which has been proposed to be generally the case among Neotropical austral migrants (Jahn and Cueto 2012, Jahn et al. 2020).

In summary, our study is the first to describe the activity patterns of a Neotropical austral migrant species during pre- and post-breeding migration, showing that elaenias are nocturnal migrants. In spite of the small sample size, our results demonstrate that accelerometer technology is a viable tool to gain deeper insights into the hourly activity profiles during migration for Neotropical austral migrants, as has already been shown in migratory passerines that breed in the Northern Hemisphere. More generally, accelerometers will allow for a more robust test of whether and how migration strategies differ across populations and migratory systems, and a deeper appreciation of how small birds are able to overcome multiple challenges on an hourly basis while on migration.

## Acknowledgments

We thank Ernesto Schajman for allowing us to work at El Principio Ranch and Feliciano Ripa for his collaboration with our research at the ranch. This work was supported by the Consejo Nacional de Investigaciones Científicas y Técnicas (CONICET-Argentina) and the Prepared for Environmental Change Initiative at Indiana University (USA).

## Supplementary material

**Figure S1.**
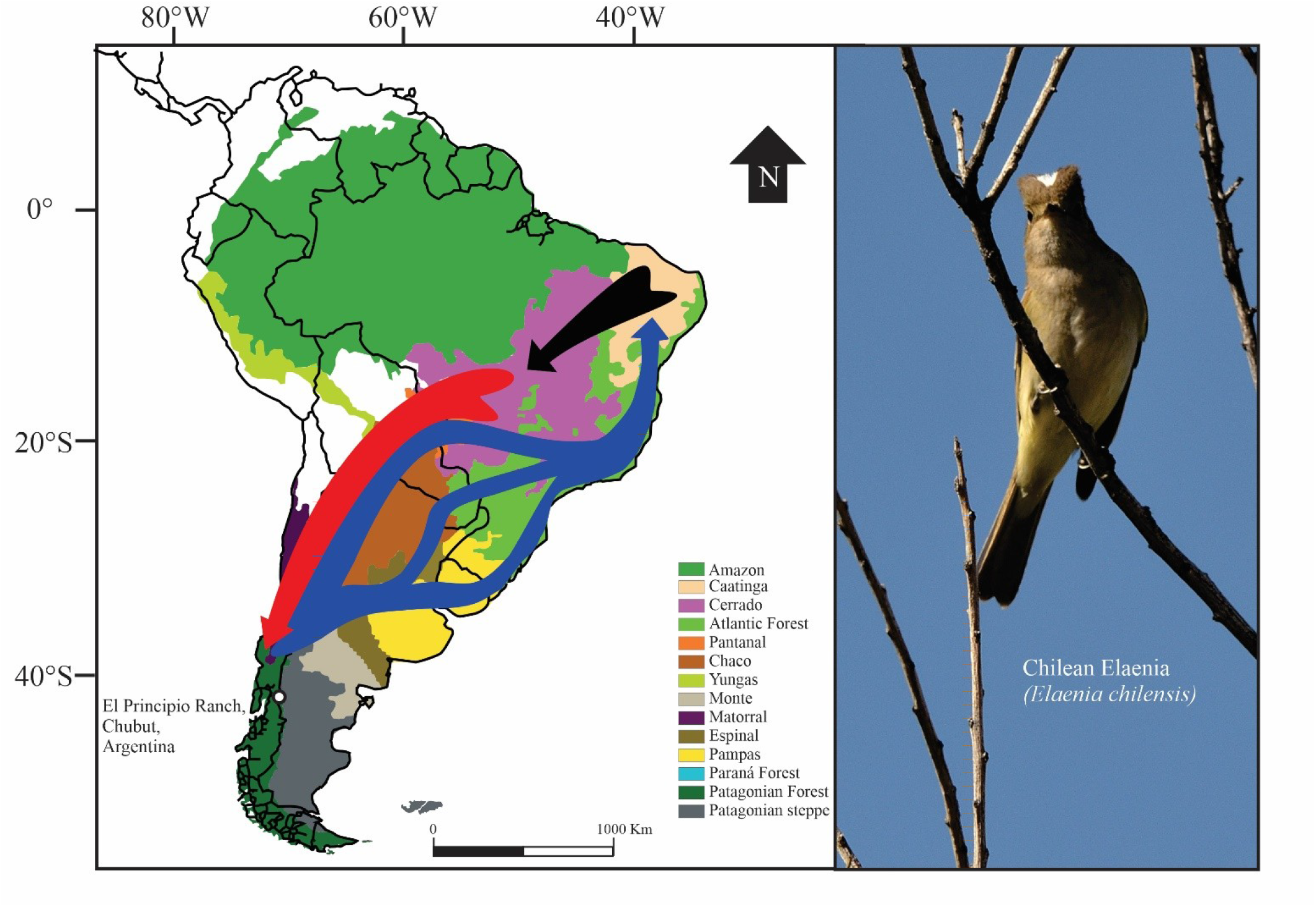
Adult Chilean Elaenia and map of South America showing study site location and migratory routes of Chilean Elaenia from previous research (Bravo et al. 2017). Blue arrows indicate post-breeding migratory routes, the black arrow indicates intra-tropical movements during the non-breeding season, and the red arrow indicates the pre-breeding migratory route.

**Table S1.**
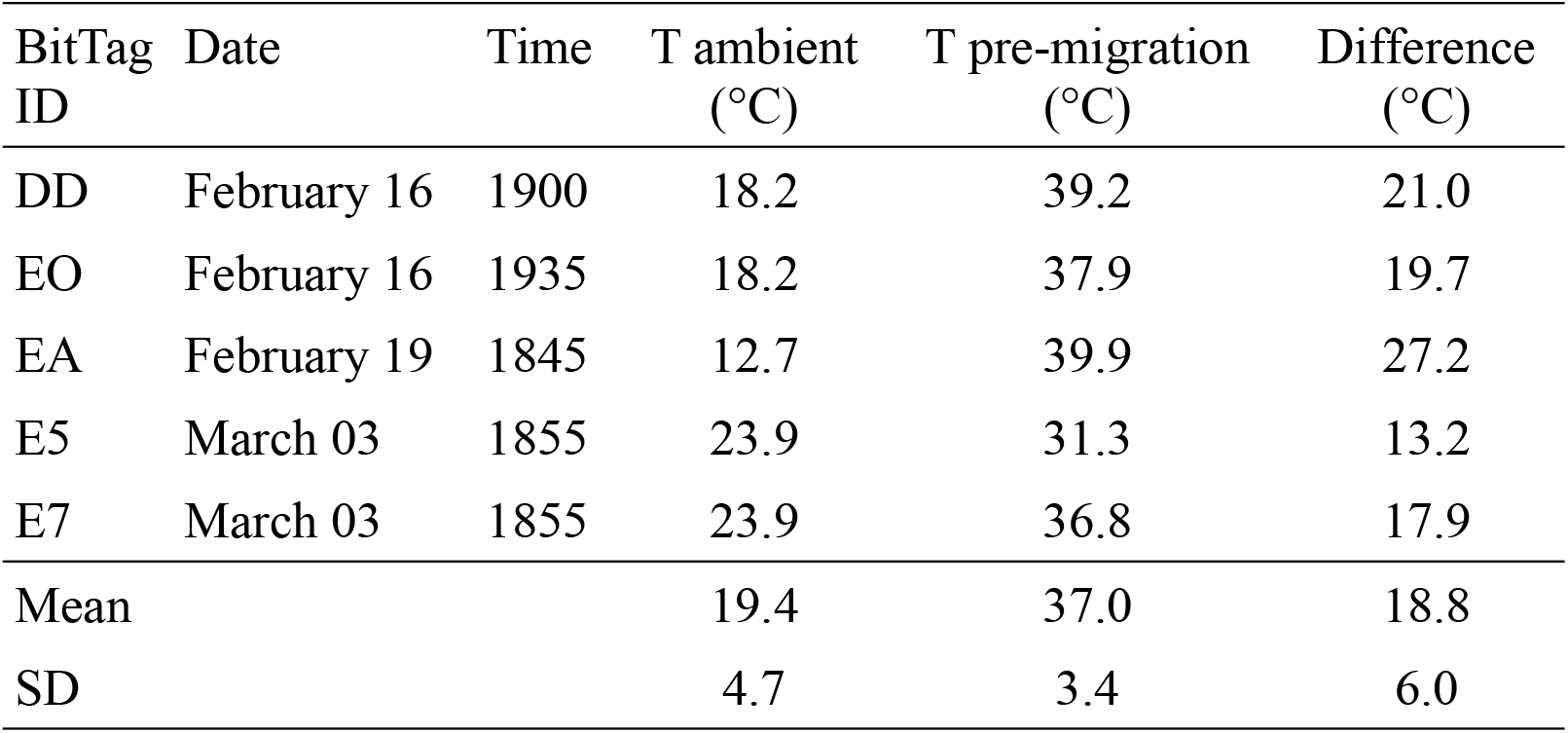
Ambient temperature at Río Percey meteorological station during the hour before the first post-migration flight and the first temperature recorded during that flight by BitTags deployed on Chilean elaenias in the forest-steppe ecotone at El Principio Ranch, Chubut Province, Argentina. Río Percey meteorological station is located 9 km NW of our study site; SD = standard deviation.

| BitTag ID | Date | Time | T ambient (°C) | T pre-migration (°C) | Difference (°C) |
| --- | --- | --- | --- | --- | --- |
| DD | February 16 | 1900 | 18.2 | 39.2 | 21.0 |
| EO | February 16 | 1935 | 18.2 | 37.9 | 19.7 |
| EA | February 19 | 1845 | 12.7 | 39.9 | 27.2 |
| E5 | March 03 | 1855 | 23.9 | 31.3 | 13.2 |
| E7 | March 03 | 1855 | 23.9 | 36.8 | 17.9 |
| Mean |  |  | 19.4 | 37.0 | 18.8 |
| SD |  |  | 4.7 | 3.4 | 6.0 |

## Notes

### Competing Interest Statement

The authors have declared no competing interest.

